# Production of novel Spike truncations in Chinese hamster ovary cells

**DOI:** 10.1101/2021.12.06.471489

**Authors:** Shiaki A. Minami, Seongwon Jung, Yihan Huang, Bradley S. Harris, Matthew W. Kenaston, Roland Faller, Somen Nandi, Karen A. McDonald, Priya S. Shah

## Abstract

SARS-CoV-2 Spike is a key protein that mediates viral entry into cells and elicits antibody responses. Its importance in infection, diagnostics, and vaccinations has created a large demand for purified Spike for clinical and research applications. Spike is difficult to express, prompting modifications to the protein and expression platforms to improve yields. Alternatively, Spike receptor binding domain (RBD) is commonly expressed with higher titers, though it has lower sensitivity in serological assays. Here, we improve transient Spike expression in Chinese hamster ovary (CHO) cells. We demonstrate that Spike titers increase significantly over the expression period, maximizing at 14 mg/L at day 7. In comparison, RBD titers peak at 54 mg/L at day 3. Next, we develop 8 Spike truncations (T1-T8) in pursuit of a truncation with high expression and antibody binding. The truncations T1 and T4 express at 130 mg/L and 73 mg/L, respectively, which are higher than our RBD titers. Purified proteins were evaluated for binding to antibodies raised against full-length Spike. T1 has similar sensitivity as Spike against a monoclonal antibody and even outperforms Spike for a polyclonal antibody. These results suggest T1 is a promising Spike alternative for use in various applications.

## Introduction

The emergence of coronavirus infectious disease 2019 (COVID-19), caused by severe acute respiratory syndrome coronavirus 2 (SARS-CoV-2), has resulted in over 250 million infections and 5 million deaths globally since November 2019. Major aspects of containing this global pandemic are surveillance (large-scale and rapid asymptomatic testing) and herd immunity (immunity achieved in a large portion of the population with protective antibodies resulting from vaccination or natural infection). Many of these containment efforts require generating large amounts of viral glycoproteins. Consequently, the COVID-19 pandemic has highlighted the critical need for rapid, scalable, and cost-effective production of recombinant glycoproteins for use as antigens in diagnostic kits, research reagents, and even the active pharmaceutical ingredient in protein-based vaccines.

For SARS-CoV-2, diagnosis and vaccination strategies involve scalable production of the Spike glycoprotein. Spike is the structural protein responsible for protecting the viral genome and for entry into cells. Spike contains the S1 and S2 domains, which mediate host receptor binding and membrane fusion, respectively (Huang et al., 2020). The receptor binding domain (RBD) of Spike lies within the S1 domain (Fig. 1A). Spike is a major antigen and the primary target for antibody binding. Consequently, immunoassays to assess immunity of individuals or a community require a SARS-CoV-2 antigen, most commonly the Spike protein. Protein-based SARS-CoV-2 vaccines also rely on delivering Spike protein with adjuvant for immunization (Heath et al., 2021).

**Figure 1.**
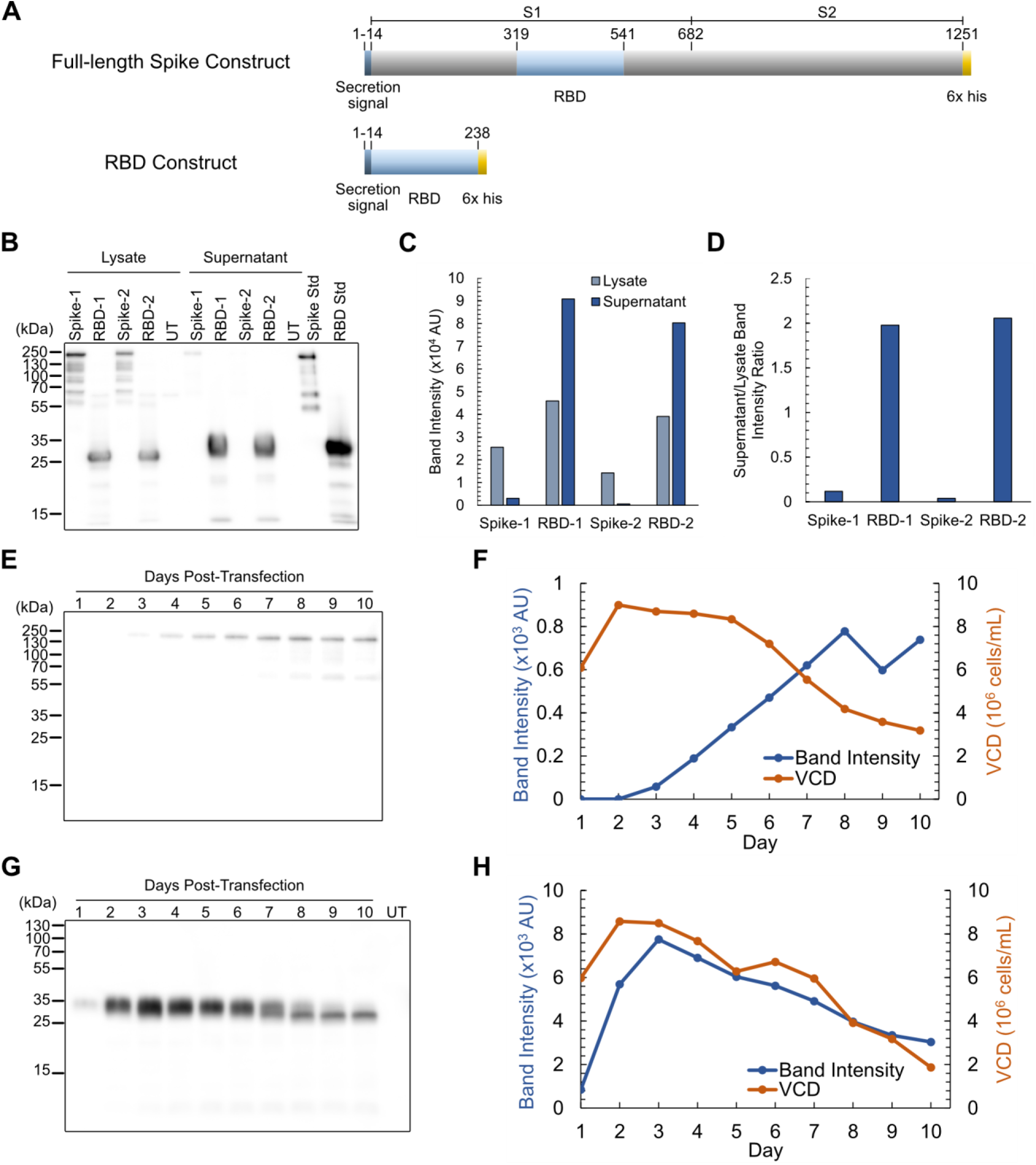
Transient Spike and RBD production in CHO cells. (**A**) Diagram of full-length Spike (1257 aa) and RBD (244 aa) constructs. Residues are labeled starting from the beginning of the secretion signal. (**B**) Western blot and (**C**) densitometry comparing two secretion signals for Spike and RBD. (**D**) Ratio of band intensities of supernatants and lysates. (**E**) Western blot and (**F**) densitometry on Spike expression time course. (**G**) Western blot and (**H**) densitometry on RBD expression time course. Abbreviations: aa (amino acids); untransfected (UT); standard (std); viable cell density (VCD).

A major limitation to scaling these approaches is generating Spike protein at high titers in a cost-effective manner. Several forms of full-length Spike have been produced in mammalian cell lines, including modifications to increase stability and expression, but titers remain at a low range of approximately 5-30 mg/L, with one report of 150 mg/L (Amanat et al., 2020; Hsieh et al., 2020; Stuible et al., 2021). A possible alternative is to express the Spike RBD, which can have expression levels of an order of magnitude higher than those of Spike, but is less sensitive than Spike in serological assays (Amanat et al., 2020). This suggests that RBD may not have the same activity as Spike for such applications. Mutational scanning has been performed on RBD, which resulted in higher expression and stability (Smaoui & Yahyaoui, 2021; Starr et al., 2020). Rational structure-based approaches have also been used to improve stability of full-length Spike (Hsieh et al., 2020). Nonetheless, identifying sequence-independent methods to increase expression is essential for comparisons with existing variants.

In this work, we transiently express Spike and RBD in Chinese hamster ovary (CHO) cells. To find high expressing and antibody binding forms of Spike, we also design and express 8 truncations of Spike, which include the RBD and additional residues. Two of these truncations express at high levels. Using simulation and experimentation, we find that one of the high-producing truncations also has more structural similarity to full-length Spike than the other and has higher binding to anti-Spike antibodies. Taken together, these truncations may provide an additional avenue for lower cost production of COVID-19 biologics with improved expression and antibody binding.

## Methods

### Plasmids

pCAGGS-Spike and pCAGGS-RBD were gifted from Florian Krammer (Amanat et al., 2020). Spike and RBD both contain an N-terminal signal peptide for secretion and a hexahistidine (6x His) tag for purification. Spike-1 and RBD-1 contain the signal sequence MFVFLVLLPLVSSQ. Spike-2 and RBD-2 contain the signal sequence MEFGLSWLFLVAILKGVQC. Spike has two stabilizing mutations (K983P and V984P), and its polybasic furin site has been removed (RRAR to R). Truncations T1-T8 were synthesized (GenScript, Piscataway, NJ) with overhangs for insertion into pCAGGS vectors (Table S1). Truncations were inserted into pCAGGS vectors via Gibson Assembly of pCAGGS-RBD digested with XbaI and XhoI. Spike truncations were designed by adding increments of approximately 50 amino acids to the N- and/or C-termini of RBD. Each truncation includes an N-terminal signal peptide and a C-terminal 6x His tag. Possible structural and binding motifs for the truncations were determined with PredictProtein (Yachdav et al., 2014). Starting and ending residues were selected to avoid interrupting major secondary structures present in Spike (Kelley et al., 2015; Meng et al., 2006; Pettersen et al., 2021).

### Cell culture and transfection

ExpiCHO-S cells (Thermo Fisher Scientific, Waltham, MA) were maintained in a 125 mL vented shake flask with 30 mL of culture in ExpiCHO Expression Medium (Thermo Fisher Scientific). Cells were cultured in a humidified incubator at 37°C and 8% CO_2_, on a 19 mm shaking diameter orbital shaker at 120 rpm (Ohaus, Parsippany, NJ).

For transfection in 125 mL shake flasks, cultures were transfected using the Expifectamine CHO Transfection Kit (Thermo Fisher Scientific), following manufacturer instructions for the Standard Protocol. For time course experiments, 0.5 mL of culture was harvested each day. Viable cell densities were measured using trypan blue and a TC20 automated cell counter (Bio-Rad, Hercules, California). Samples were harvested by centrifuging at 300 rcf for 5 min and collecting the supernatant. For samples to undergo purification, entire cultures were centrifuged at 4,000 rcf for 20 minutes at 4°C and filtered through 0.22 µm filters. For transfection in 2.0 mL 96-well deep well blocks (Genesee Scientific, El Cajon, CA), 0.8 mL of cells at 6 × 10^6^ cells/mL were plated on the day of transfection. Cells were cultured on a 3 mm shaking diameter orbital shaker at 900 rpm (Benchmark Scientific, Sayreville, NJ) and transfected according to manufacturer instructions. Samples were harvested 5 days post-transfection by centrifuging the cultures at 300 rcf for 5 minutes and collecting the supernatant.

### Protein purification and concentration

Filtered samples were column purified using an AKTA Pure fast protein liquid chromatography (FPLC) system with a 5 mL prepacked Ni Sepharose HP column (Cytiva, Marlborough, MA), using imidazole to elute the proteins (Esposito et al., 2020). Samples were loaded onto the column at a flow rate of 5 mL/min, the resin was washed for 10 column volumes (CV), and proteins were eluted using imidazole. Detailed procedures are available in Supporting Information. Purified proteins were dialyzed against phosphate-buffered saline (PBS) using dialysis cassettes at 4°C (Thermo Fisher Scientific). Spike was dialyzed with a 20 kDa molecular weight cutoff (MWCO) membrane. RBD, T1, and T4 were dialyzed with 10 kDa MWCO membranes. Dialyzed samples were concentrated using centrifugal filter units (Millipore Sigma, Burlington, MA) at 4,000 rcf for 20 minutes at 4°C. Spike was concentrated using centrifugal filter units with a MWCO of 30 kDa. RBD, T1, and T4 were concentrated with 10 kDa MWCO centrifugal filter units.

### SDS-PAGE and western blot

Samples from time course experiments and the truncation screening were prepared for sodium dodecyl-sulfate-polyacrylamide gel electrophoresis (SDS-PAGE) by adding 12 µL of NuPAGE LDS Sample Buffer (Thermo Fisher Scientific) and 3 µL of tris(2-carboxyethyl)phosphine (Thermo Fisher Scientific) to 30 µL of sample. Mixtures were heated at 95°C for 10 minutes and 10 µL of samples were loaded into gels cast in-house, with a 12% acrylamide resolving layer and 4% acrylamide stacking layer. Samples were run through the gel for 15 minutes at 115 V, then 50 minutes at 150 V. Proteins were transferred onto polyvinylidene difluoride membranes in a wet sandwich and membranes were blocked using 5% non-fat milk. Membranes were stained overnight at 4°C with a 1:1000 diluted mouse anti-his primary antibody (MCA1396, RRID:AB_322084, Bio-Rad) and then for 1 hour at room temperature with a 1:4000 diluted rabbit anti-mouse HRP secondary antibody (SouthernBiotech Cat# 6170-05, RRID:AB_2796243, Birmingham, AL). Membranes were developed using Pierce ECL Western Blotting Substrate (Thermo Fisher Scientific) and imaged using an Amersham Imager 600 (Cytiva).

Purified samples were analyzed by SDS-PAGE with a method previously described (Xiong et al., 2019). Images of the gels were taken using a ChemiDoc Imaging system (Bio-Rad), and proteins were transferred onto nitrocellulose membranes using Trans-Blot Turbo Packs (Bio-Rad) and Trans-Blot Turbo System (Bio-Rad). Membranes were blocked overnight in 1% casein, stained with 1:1000 diluted mouse anti-his primary antibody and stained with 1:4000 diluted rabbit anti-mouse secondary antibody. The chemiluminescent reactions were performed using Clarity ECL Substrate (Bio-Rad).

Concentrations for purified proteins were estimated using a combination of ELISA, Bradford Assay, and scanning densitometry on SDS-PAGE gels. Spike and RBD concentrations were first calculated using sandwich ELISA. Purified T1 and T4 concentrations were determined using Bradford Assay since ELISA standard curves could not be generated for these novel protein truncations. Next, 1 µg of proteins, as determined by the two methods, were loaded into each lane of a 4%-20% gradient stain-free gel (Bio-Rad). Dilutions of RBD standard from 1.5 µg to 0.5 µg were also loaded into the gel. Samples were run at 200 V for 36 minutes and imaged using a ChemiDoc imaging system (Bio-Rad). A standard curve was generated via densitometry through ImageJ, and primary band intensities for the samples were interpolated to quantify concentrations.

### Enzyme-linked immunosorbent assay (ELISA)

Sandwich ELISAs were performed to quantify purified Spike and RBD and crude supernatants. 1:1000 mouse anti-his capture antibody in PBS was coated onto Immulon 2 HB 96-well plates (Thermo Fisher Scientific) at 4°C overnight. Plates were blocked with 200 µL/well PBS with 3% BSA for 30 minutes. Plates were loaded with serial dilutions of purified protein samples or crude supernatants. Plates were incubated with 1:1000 rabbit anti-RBD primary antibody (Sino Biological Cat# 40592-R001, RRID:AB_2857936, Wayne, PA), then 1:6000 or 1:4000 goat anti-rabbit, HRP secondary antibody (SouthernBiotech Cat# 4030-05, RRID:AB_2687483) in PBS with 1% BSA for purified or crude proteins, respectively. Plates were developed with 1-step Turbo TMB-ELISA Substrate Solution (Thermo Fisher Scientific) and 2N HCl. Absorbance at 450 nm was measured using a Spectramax 250 spectrophotometer (Molecular Devices, San Jose, CA). Plates were washed 3 times with 200 µL/well PBS with 0.05% Tween20 (PBS-T) between each step and incubations were performed using volumes of 100 µL/well for 1 hour at room temperature unless specified otherwise. Standard curves for quantifying Spike and RBD were generated using serial dilutions of Sf9 insect Spike (NR-52308, BEI Resources, Manassas, VA) and HEK293F human RBD (NR-52366, BEI Resources), respectively.

Indirect ELISAs were performed to assess the sensitivities of CHO-expressed proteins to a human anti-Spike monoclonal antibody CR3022 (NR-52392, BEI Resources, RRID:AB_2848080) and a rabbit anti-Spike polyclonal antibody (PAb, eEnzyme, SCV2-S-100, RRID:AB_2893135, Gaithersburg, MD). For CR3022, antigens were first coated onto plates at 4°C overnight. After blocking, serial dilutions of CR3022 in PBS with 1% BSA were loaded from 100 ng/well. Plates were loaded with 100 µL/well goat anti-hIgG, HRP secondary antibody at 1:4000 in PBS containing 1% BSA. For the PAb, 3-fold serial dilutions starting at 400 ng/well of rabbit anti-Spike primary antibody were used (PAb, SCV2-S-100, eEnzyme), and a 1:4000 goat anti-rabbit IgG, HRP secondary antibody was used instead.

### Bradford assay

Bradford assays were performed to quantify the concentration of total soluble protein (TSP) by using a protein assay dye reagent (Bio-Rad). For each BSA standard, sample, and diluted sample, 10 µL/well of sample and 190 µL/well of Bradford dye were loaded into 96-well plates. After incubating for 10 minutes at room temperature, the absorbances of standards and samples were measured at 450 nm and 590 nm (Ernst & Zor, 2010), using a Spectramax M4 spectrophotometer (Molecular Devices). Standard curves for quantifying samples were generated by using serial dilutions of BSA from 0-0.5 mg/mL with 0.05 mg/mL steps.

### Liquid chromatography-tandem mass spectrometry (LC-MS/MS)

Sequences of purified T1 and T4 were obtained via LC-MS/MS. 10 µg of T1 and 20 µg of T4 were run on a 4%-20% gradient SDS-PAGE gel. Bands were extracted and submitted to the Genome Center Proteomics Core at the University of California, Davis. Briefly, proteins were digested with tryspin and analyzed on a Dionex UltiMate 3000 RSLC system (Thermo Fisher Scientific) using a PepSep (PepSep, Denmark) ReproSil 8 cm 150 µm I.D. C18 column with 1.5 μm particle size (120 Å pores). Mass spectra analysis is described in Supporting Information. Searches were conducted against the known sequences of T1 and T4, and alignments were performed using Multiple Alignment using Fast Fourier Transform (Katoh et al., 2019).

### Circular dichroism (CD)

Concentrated samples were prepared for CD analysis by diluting 150 µg of protein in 50% PBS and 50% CD buffer (25mM of phosphate and 40mM of NaF). Single spectrum data were obtained using a Jasco J-715 CD spectrometer (Jasco, Easton, MD). Data were analyzed using BeStSel (Micsonai et al., 2015). Spectra of buffer were subtracted before analysis. To obtain secondary structure data for the PDB Spike structure, the PDB file 6VXX was analyzed using the STRIDE server (Frishman & Argos, 1995).

### Simulations

Starting configurations for molecular dynamics (MD) simulations were obtained by trimming the full Spike protein structure obtained from the protein data bank (6VXX). Structures were reduced to a single monomer and cut at the amino acid sequences corresponding to RBD, T1, T3, and T4. 6x His tags were added using modeller (Webb & Sali, 2016), which modifies amino acid sequences of proteins. The new His-tagged structures were prepared and had glycans attached using Glycam (Woods, 2005). The N-glycosylation sites of RBD and the RBD portion of all truncations had the glycoform FA2 attached. T1 contained no additional N-glycosylation sites, T3 contained an additional FA3 glycoform, and T4 contained an additional M5 glycoform. Amber ff14SB and Glycam06 forcefields (Kirschner et al., 2008; Maier et al., 2015) were used and generated using acpype.py following the method shown previously (Bernardi et al., 2017, 2019). Simulations were conducted using the Gromacs 2019.1 suite with similar energy minimization procedure as in previous simulations (Abraham et al., 2015; Pronk et al., 2013; Van Der Spoel et al., 2005) including ones involving glycosylated RBD (Bernardi et al., 2017; Huang et al., 2021). Simulation runs after equilibration were carried out for 100 ns.

## Results

### Expression and purification of Spike and RBD

We first compared the expression of Spike and RBD in ExpiCHO-S cells transfected in 96-well format. Spike and RBD were expressed with N-terminal secretion signals and C-terminal 6x His tags for downstream purification (Fig. 1A). We also replaced the previously tested secretion signal with an alternative secretion signal to determine whether it affects expression and secretion of Spike and RBD (Spike-1, Spike-2, RBD-1, and RBD-2, Fig. 1B).

Relative amounts of protein in the supernatant and cell lysate were determined by western blot 5 days post-transfection (Fig. 1B and 1C). Spike had significantly lower expression than RBD and was particularly less abundant in the supernatant. Comparison of ratios of supernatants over lysates also showed that Spike is significantly retained in the cells compared to RBD (Fig. 1D). Both the expression and supernatant/lysate ratio remained the same for Spike and RBD with either signal sequence, indicating that low expression and high retention of Spike in the cells may be due to the protein sequence itself, and not a consequence of the tag used. All following experiments were performed with Spike-1 and RBD-1, hereafter referred to as Spike and RBD, respectively.

Next, cultures were scaled up to 25 mL and a time course experiment was performed to determine the optimal harvest time for maximum titers. Cells were transfected with plasmids encoding Spike and RBD, and a sample of the supernatant was collected every 24 hours over 10 days. Western blots were performed on the supernatants and band intensities were plotted over time (Fig. 1E-H). Spike concentration in the supernatant increased steadily until 7 days post-transfection, after which time it remained stable (Fig. 1E and 1F). In comparison, RBD concentration in the supernatant peaked at day 3, then decreased (Fig. 1G and 1H). From these results, Spike and RBD harvests were determined to be 7 and 3 days post-transfection, respectively.

To produce large quantities of Spike and RBD for purification and downstream analysis, 150 mL of supernatants were prepared from pooled 25 mL cultures. Crude titers of Spike and RBD were measured using sandwich ELISAs on filtered crude, yielding 14 mg/L and 54 mg/L, respectively (Fig. S1). The crude supernatants were purified through FPLC (Fig. 2 and S2). For purification of Spike, SDS-PAGE revealed bands in elution fractions E3, E4, and E5 (Fig. 2A), which were confirmed by western blot to include immunoreactive bands consistent with full-length Spike (Fig. 2B). For RBD, SDS-PAGE (Fig. 2C) and western blot (Fig. 2D) showed bands in elution fractions E3, E4, and E7. FPLC-purified samples were dialyzed using PBS and concentrated through centrifugal filter units. E3 fractions of both proteins were used for subsequent experiments.

**Figure 2.**
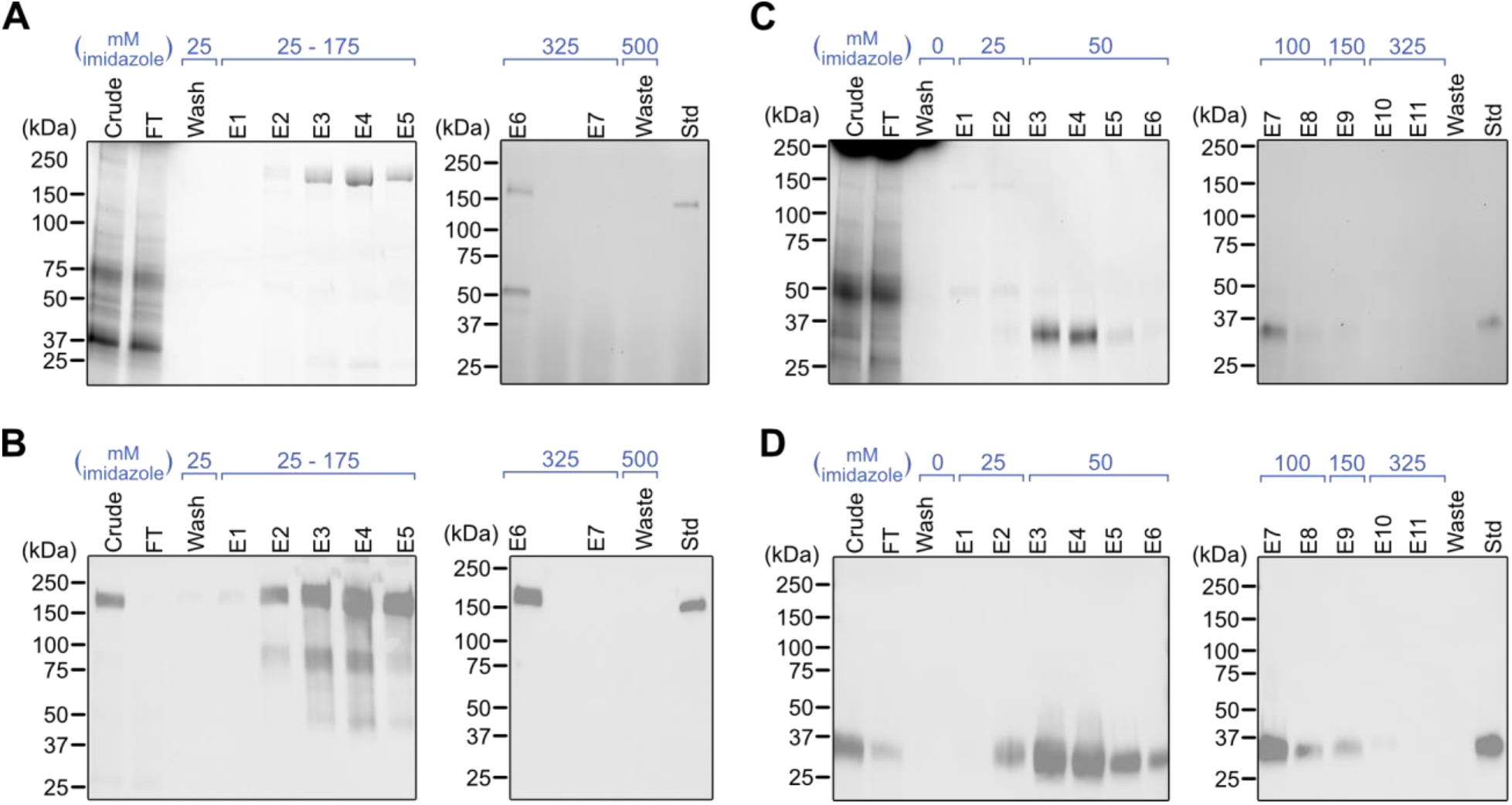
Purification of Spike and RBD. (**A**) SDS-PAGE and (**B**) western blot on Spike fractions. (**C**) SDS-PAGE and (**D**) western blot on RBD fractions. FT samples were collected during sample loading onto the column. FT and wash samples were pooled from multiple fractions at equal volumes. Abbreviations: flow-through (FT); elution (E); standard (std).

### Novel truncations to improve protein titers

Given the difficulty in expressing full-length Spike but its higher sensitivity in serological assays (Amanat et al.), we sought to determine whether other truncations that are larger than the RBD could be highly expressed and maintain high sensitivity. We developed eight truncations, T1-T8, by adding increments of approximately 50 amino acids on the N- or C-terminal ends of RBD (Fig. 3A). We also expressed the full S1 subunit of Spike. Secretion signals and 6x His tags were added to N- and C-termini, respectively.

**Figure 3.**
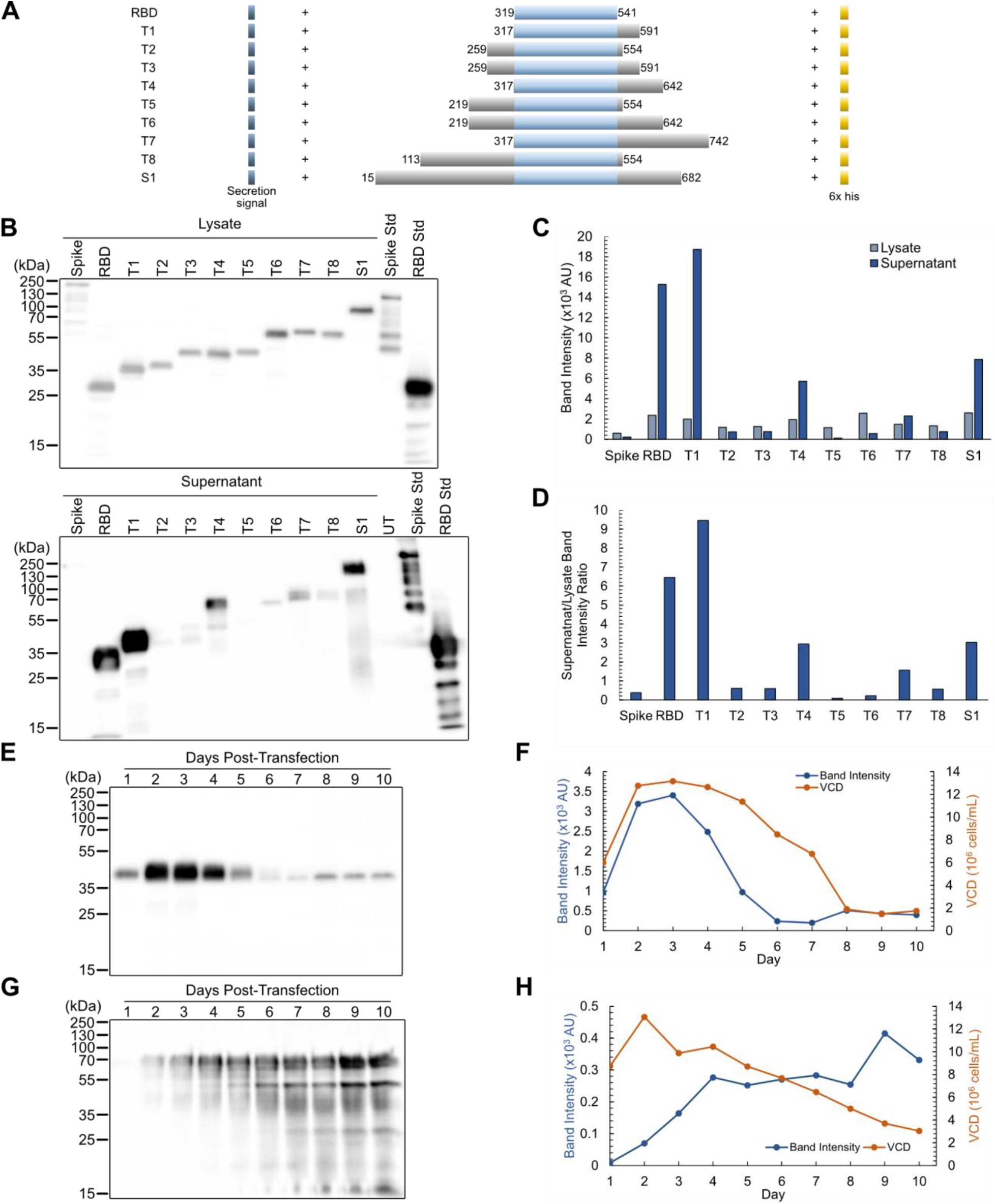
Expression of Spike truncations. (**A**) Construct diagram of Spike truncations. Residue numbers are relative to the position within full-length Spike, including the secretion tag. (**B**) Western blot and (**C**) densitometry on truncations for lysates and supernatants. (**D**) Ratio of band intensities of supernatants over lysates. (**E**) Western blot and (**F**) densitometry of T1 expression over time. (**G**) Western blot and (**H**) densitometry of T4 expression over time. Abbreviations: untransfected (UT); standard (std); viable cell density (VCD).

We first screened the truncations for expression levels. Cells were transfected in 96-well format, and lysates and supernatants were analyzed by western blot 5 days post-transfection (Fig. 3B). T1 and T4 had the highest expression, as well as the highest supernatant/lysate signals (Fig. 3C and 3D). In particular, T1 had even higher expression and relative secretion than RBD did. Given that S1 has previously been studied (Ren et al., 2020), T1 and T4 were selected for scaleup and purification. To determine optimal harvest dates for T1 and T4, and expression time course was performed as previously described for Spike and RBD. For T1, western blot and densitometry (Fig. 3E and 3F) showed that expression peaked 3 days post-transfection with a single band. For T4, expression peaked after 4 days post-transfection, with degradation bands also increasing after this point (Fig 3G and 3H). Therefore, T1 and T4 harvest dates were determined to be 3 and 4 days post-transfection, respectively.

For purification, transfections with T1 and T4 were performed in 150 mL cultures and samples were purified through FPLC (Fig. 4 and S3). SDS-PAGE revealed two bands for T1, one of which was determined to be T1 through western blot (Fig. 4A and 4B). To remove the impurity, T1 was dialyzed and repurified. For T4, three bands were detected: one band at 70 kDa, one band at 50 kDa, and one band at slightly below 50 kDa (Fig. 4C). Western blot detected bands in both the 70 kDa and 50 kDa regions (Fig. 4D), indicating two forms of T4. The impurity was removed through dialysis and repurification. Pure T1 and T4 were obtained in this manner and prepared for further characterization through dialysis and spin column concentration. The bands detected for T1 and T4 through western blot were cut out of an SDS-PAGE gel and submitted for proteomic analysis. Coverage for T1 and T4 were high (Fig. S4), and the two bands of T4 on the SDS-PAGE gel suggest different post-translational modifications. Quantifying crude titers, we found that T1 and T4 expressed at 130 mg/L and 73 mg/L, respectively, which are higher than our RBD titers (Fig. S5). For further characterization and binding assays, concentrations of purified samples were determined using Sandwich ELISAs and SDS-PAGE densitometry (Fig. S6).

**Figure 4.**
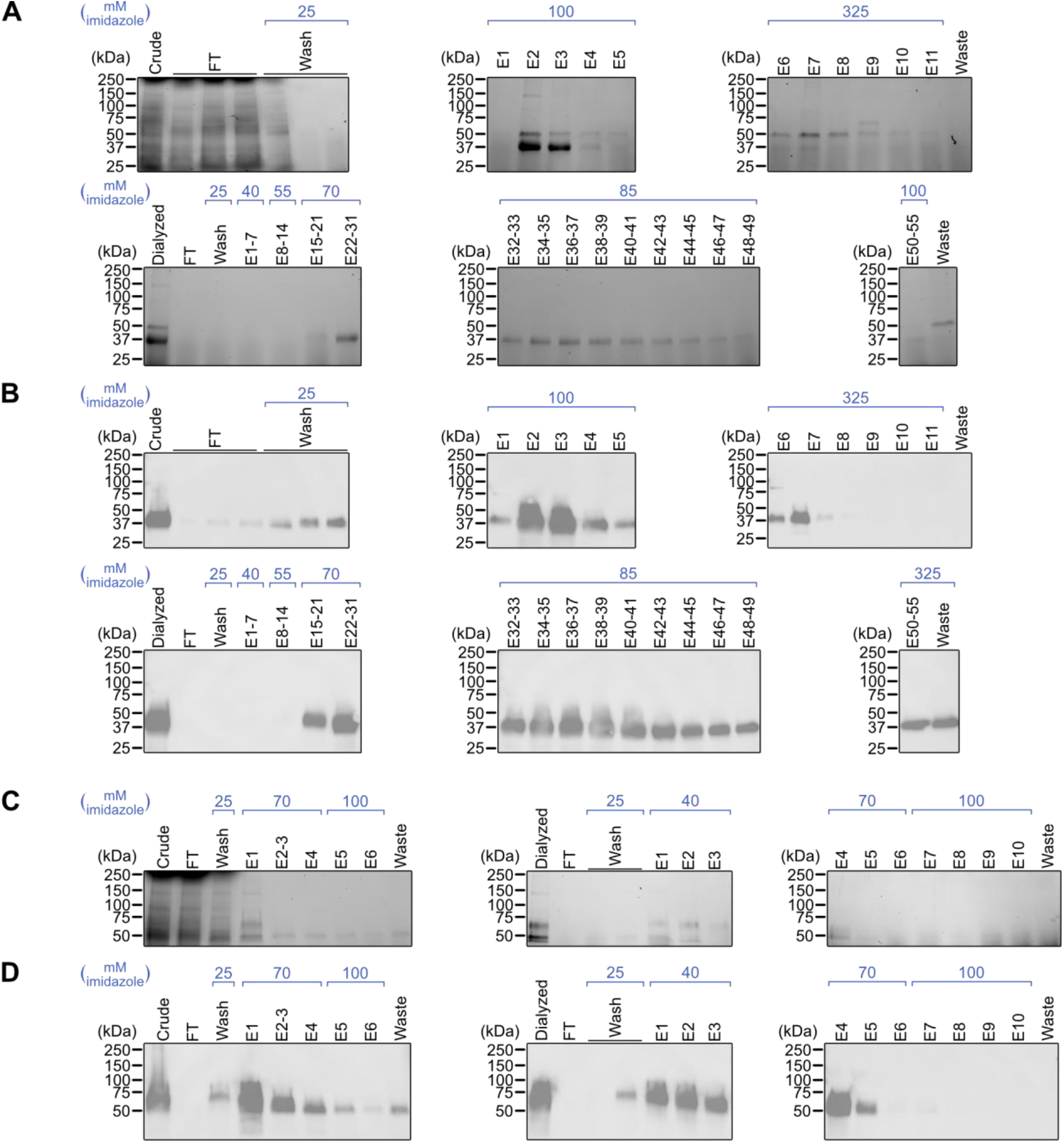
Purification of T1 and T4. (**A**) SDS-PAGE and (**B**) western blot on T1 fractions from crude purification and repurification. (**C**) SDS-PAGE and (**D**) western blot on T4 fractions from crude purification and repurification. FT and wash samples were pooled from multiple fractions at equal volumes. Abbreviations: flow-through (FT); elution (E).

### Binding sensitivities against antibodies

The activities of the CHO-expressed proteins were evaluated via indirect ELISAs with antibodies raised against full-length Spike. First, the monoclonal antibody CR3022 was tested, which binds to the receptor binding domain of Spike (Yuan et al., 2020). Serial dilutions of CR3022 were incubated with Spike, RBD, T1, and T4 (Fig. 5A). Binding sensitivities were compared by taking the areas under the curves (Fig. 5B). To compare with another recombinant source of Spike, Sf9 insect cell-expressed Spike was also used in the assays.

**Figure 5.**
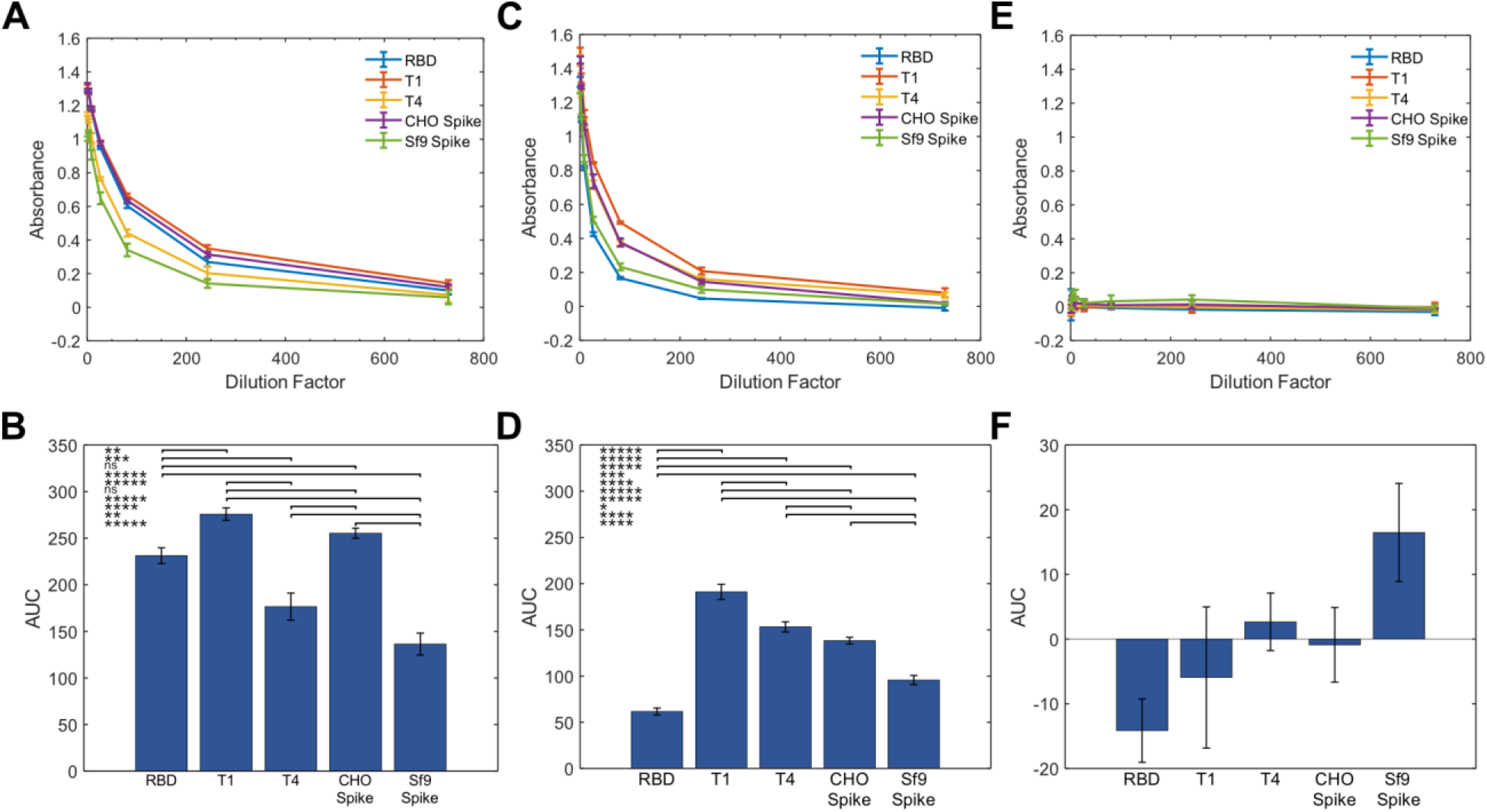
Binding assays of Spike truncations with anti-Spike antibodies. (**A**) Absorbance as a function of dilution factor of CR3022. (**B**) AUC calculated from (A). (**C**) Absorbance as a function of dilution factor of an anti-Spike PAb. (**D**) AUC calculated from (C). (**E**) Absorbance against dilution factor of a rabbit normal IgG antibody. (**F**) AUC calculated from (E). Error bars represent ± SD of technical triplicates. p-values were calculated using a one-way ANOVA followed by Tukey’s Test. * indicates p < 5×10^−2^, ** p < 5×10^−3^, *** p < 5×10^−4^, **** p < 5×10^−5^, and ***** p < 5×10^−6^. Abbreviations: area under curve (AUC); ns (not significant).

CHO-expressed Spike had higher binding to CR3022 than Sf9-expressed Spike. This may be due to differences in folding or glycosylation between the CHO- and insect-expressed proteins. Among the CHO-expressed proteins, T1 had higher binding to CR3022 than RBD did and is comparable to the performance of Spike. T4 had lower signal but still outperformed Sf9-expressed Spike. Next, serial dilutions of a polyclonal antibody raised against full-length Spike were tested (Fig. 5C and 5D). Given that PAbs may recognize multiple binding epitopes in a protein, larger forms of Spike were expected to have higher performance. Strikingly, however, T1 and T4 had very high signal across dilutions of the antibody, and T1 outperformed full-length CHO-expressed Spike. The increased sensitivities were not due to non-specific binding of T1 and T4 to rabbit antibodies, since a control rabbit IgG did not produce significant signal (Fig. 5E and 5F).

### Structural characterization of truncations

To determine whether structural similarities are maintained between the truncations and the relevant regions of Spike, structures of T1 and T4 were predicted using MD. Snapshots of simulated structures of RBD, T1, and T4 at 0 ns and 100 ns simulation times are shown (Fig. 6A-D). The RBD portion of all structures remained stable during this time. T1 and T4 showed similarly stable secondary structures in the additional residues at the bottom of the structure. The more flexible turn features curled in and stabilized over the course of the trajectory. To quantify this behavior, the root mean squared deviation (RMSD) of the whole structures and RBD subdomains for each truncation were evaluated (Fig. 6E and 6F). These RMSD plots show deviation relative to initial structures and provide further evidence that the RBD subdomains are stable or reach a stable structure early in the simulation trajectory. The T1 and T4 RMSD plots show more conformational change, likely due to the flexibility of the turn features observed in the snapshots. An interesting observation is that the RBD with a 6x His tag appeared to be more stable compared to the RBD without the tag. Based on this RMSD data, T1 appeared to stabilize the RBD in line with this 6x His tag, while the T4 structure aligned more closely to the RBD without a 6x His tag.

**Figure 6.**
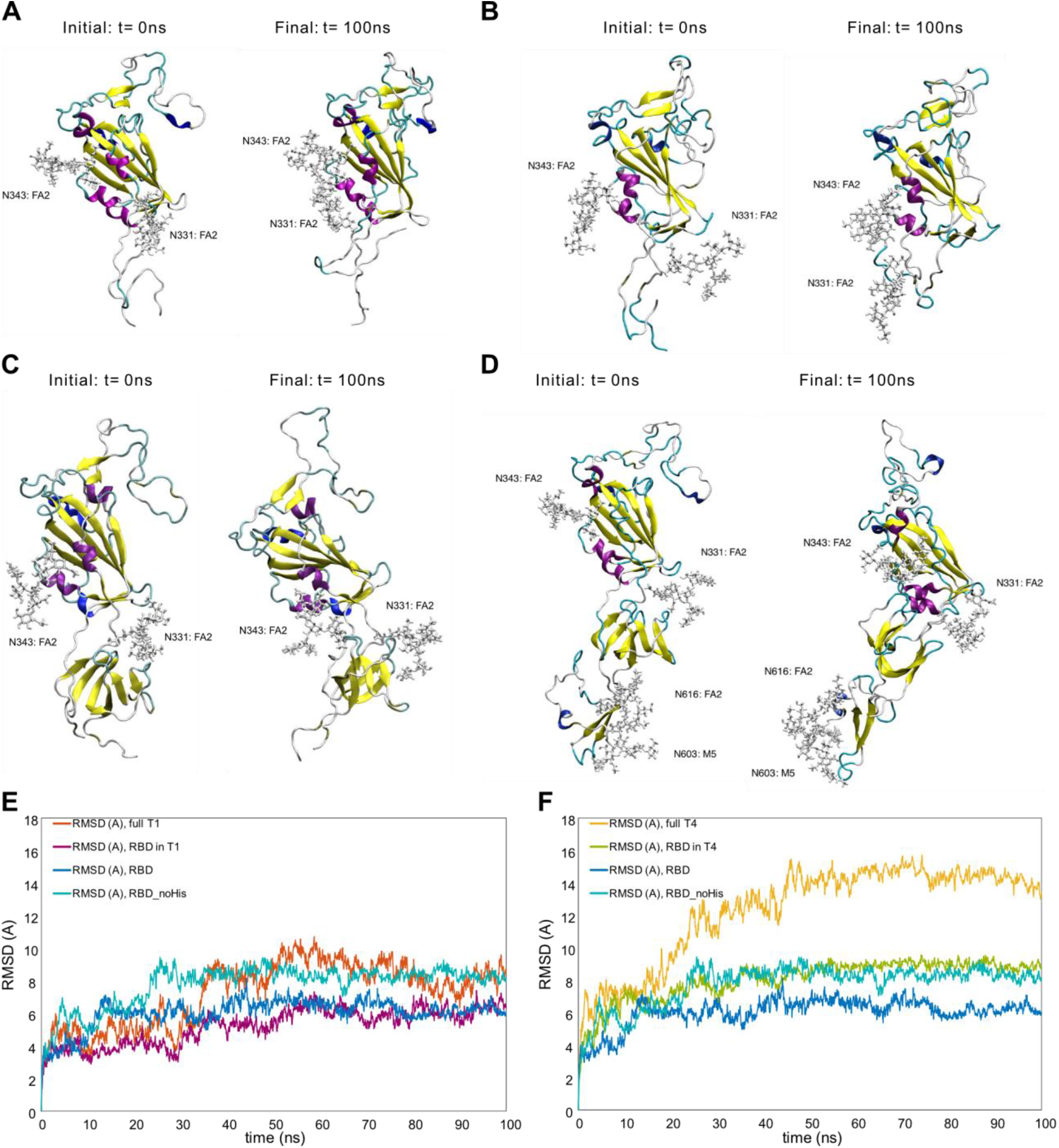
MD structural stability snapshots and analysis. MD snapshots are visualized for (**A**) RBD, (**B**) RBD without the 6x His tag (RBD_noHis), (**C**) T1, and (**D**) T4 at 0 ns and 100 ns. Backbone RMSD profiles of (**E**) full T1 and T1 RBD subdomain and (**F**) full T4 and T4 RBD subdomains are compared against RBD with and without His tag referenced to initial configurations.

We hypothesize that some truncations did not express well because of structural differences. To explore this idea, we compared T1 and T3, which only differ by ∼50 amino acids at the N-terminus but had vastly different expression (Fig. 3B and Fig. 3C). MD was used to determine whether structural differences may have caused the discrepancy in expression. RMSD analysis showed that T3 had much higher RMSD compared to T1 (Fig. S7). Visualization of T3 revealed that the difference in RMSD was due to the additional FA3 glycan binding to its own RBD, which could contribute to low expression. Removal of the FA3 glycan from T3 resulted in a secondary structure that matched more closely to T1 and a more stable RBD within T3 (Figure S7). It is possible that other truncations also had incorrect folding.

Experimentally, secondary structure compositions of CHO-expressed Spike, RBD, T1, and T4 were obtained using circular dichroism (CD). Δε values were obtained, which is a measure of the difference in absorbance of left- and right-circularly polarized light. Using the BeStSel server, Δε as a function of wavelength was analyzed to predict the secondary structure compositions. The distributions of observed secondary structures were similar for most proteins (Fig. 7A and S8). CHO Spike and Sf9 Spike had very similar compositions, suggesting high structural similarity. RBD and T1 also had similar compositions. T4 was slightly dissimilar, with low beta sheet content compared to other proteins. CD-analyzed proteins were also compared to a structure of Spike determined through cryo-electron microscopy (PDB 6VXX, Walls et al., 2020). 6VXX had similar alpha helix and beta sheet content as CHO and Sf9 Spike, but had much higher turn content and lower “other” content, which includes coils, bends, irregular loops, β-bridges, 3_10_ helices, and π-helices.

**Figure 7.**
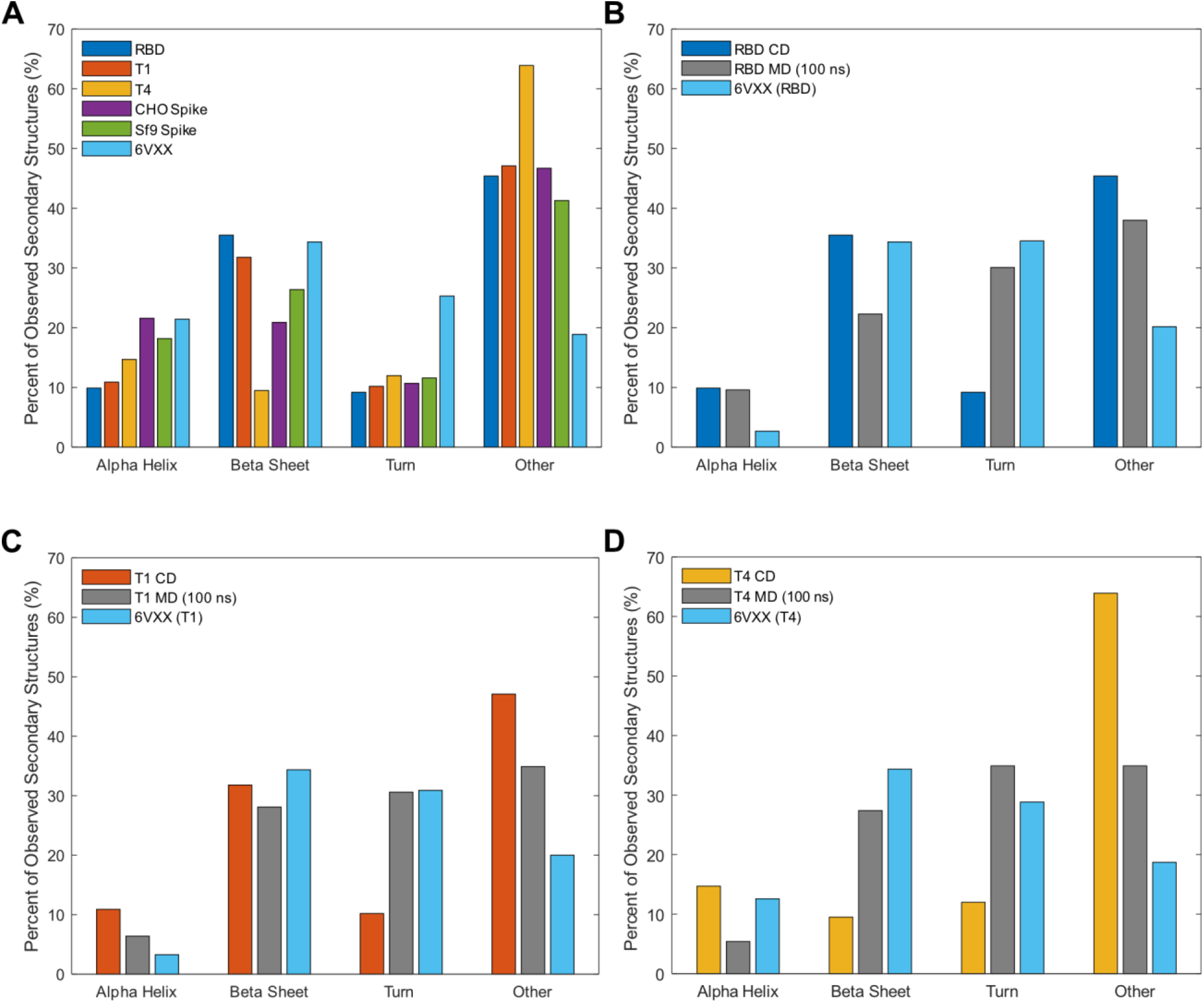
Structural composition of Spike and truncations produced in CHO cells. (**A**) Overall secondary structure compositions of CHO-expressed proteins and 6VXX. (**B**) Comparison of RBD MD to CD data and the RBD region of 6VXX. (**C**) Comparison of T1 MD to CD data and the T1 region of 6VXX. (**D**) Comparison of T4 MD to CD data and the T4 region of 6VXX. MD structural data represent proteins including the 6x His tags with the final structural compositions at 100 ns.

CD-derived structural information was also compared with MD secondary structures for RBD, T1, and T4 determined using DSSP (Kabsch & Sander, 1983) (Fig. 7B-D). Truncated 6VXX structures containing the relevant residues were also included, which represent structural composition had truncating Spike not resulted in any structural changes. For all three proteins, 6VXX and MD structures had high similarity, with CD-derived structures having lower turn content. Overall, MD and CD results suggest that T1 and T4 retain accurate RBD structure, and consistent with their high sensitivities in the ELISAs against anti-Spike antibodies.

## Discussion

Production of Spike fragments is important for its use in diagnostics, protein subunit vaccines, and research. In addition, high affinities of the Spike fragments are critical in these applications. Several approaches have been used to increase Spike yields, including stabilizing mutations (Hsieh et al., 2020), comparison of different cell lines (Stuible et al., 2021), and optimization of production conditions, such as temperature shifts (Johari et al., 2021). Here, we expressed full-length Spike and RBD transiently in CHO cells to determine the intracellular and extracellular production kinetics. In addition, we developed 8 truncations in pursuit of a truncation which exhibits both high expression and binding to antibodies.

The regions of Spike that cause lower expression and higher sensitivity compared to RBD are not known, but the initial screen of the truncations showed that T1 is highly expressed and secreted compared to other truncations, with T4 following at much lower titers (Fig. 3C and 3D). This suggests that residues downstream of T1 may be contributing to decreased titers. Comparing T1 to T2 and T3, residues upstream of T1 also appear to decrease titers. The additional residues in T2, T3, and T4 contain predicted glycosylation sites, which may introduce avenues for protein retention such as incomplete glycosylation. This is supported by the lysate proteins running at their expected molecular weights and the supernatant proteins much higher, though protein size did not appear to correlate with relative retention in the cell (Fig. 3B). In contrast, T1 only contains the same glycosylation sites as RBD and was found in the crude at much higher titers compared to other truncations. Interestingly, the MD simulations of T3 suggest that lower stability may result from unexpected intramolecular glycan-protein interactions for fully glycosylated truncations.

In the ELISA sensitivity assays for CR3022 and the PAb, CHO-expressed Spike has higher AUCs for both antibodies compared to Sf9-expressed Spike (Fig. 5). The discrepancy may be due to different glycosylation profiles, which would be consistent with the idea that CHO-expressed proteins tend to have more human-like glycosylation patterns (Esko & Stanley, 2015). We also found that CHO Spike produces higher signal than RBD when probed with the PAb, consistent with results from serological assays (Amanat, 2020). This was also expected because polyclonal antibodies target multiple epitopes, and full-length Spike may contain more binding epitopes than RBD. Surprisingly, T1 and T4 have higher sensitivities to the PAb, outperforming full-length Spike. One possibility is that T1 and T4 contain an additional epitope, not present on RBD, that has high affinity but is sterically hindered when additional residues are present. This may also be the reason for the higher performance of T1 over T4. Visualization of binding through cryo-electron microscopy and analysis of binding kinetics and thermodynamics through methods such as biolayer interferometry and steered MD may elucidate the reason for their high affinities.

## Conclusions

We expressed SARS-CoV-2 Spike and RBD in CHO cells and optimized harvest dates. Additionally, we expressed 8 new truncations and found that T1 and T4 have high expression and secretion, where T1 has even higher expression than RBD. T1 and T4 also have higher binding sensitivity to a PAb compared to Spike. Overall, T1 had the highest performance in all expression and binding experiments conducted in this work. Its high expression and sensitivity suggest T1 may be a promising Spike alternative in research and clinical applications. Further work is needed to understand why T1 has higher affinity to antibodies and whether the higher affinity translates to assays with convalescent sera.

## Acknowledgments

We would like to acknowledge the Protein Structure and Dynamics Core, Biochemistry and Molecular Medicine, University of California, Davis for obtaining CD spectral data of purified proteins. We would like to acknowledge the Proteomics Core Facility, University of California, Davis for proteomics analysis on T1 and T4. LC/MS was supported by NIH grant 1S10OD026918-01A1. We thank Nitin Beesabathuni and Oanh Pham for scientific and editorial feedback on the manuscript. Computer simulations were performed on the hpc1/hpc2 servers at UC Davis. YH and RF were partially supported by the National Science Foundation under grant no. CBET 1911267. BSH was partially supported by LLNL’s LDRD program, under the auspices of the U.S. Department of Energy by Lawrence Livermore National Laboratory under Contract DE-AC52-07NA27344.

