## Supporting Information for "Production of novel Spike truncations in Chinese hamster ovary cells"

Shiaki A. Minami[1], Seongwon Jung[1], Yihan Huang[2,#], Bradley S. Harris[1,#], Matthew W. Kenaston[3], Roland Faller[1], Somen Nandi[1,4], Karen A. McDonald[1,4], Priya S. Shah[1,3,*]
1 Department of Chemical Engineering
2 Department of Materials Science and Engineering
3 Department of Microbiology & Molecular Genetics
4 Global HealthShare Initiative
University of California, Davis

### These authors contributed equally

#### Supporting Information

**Table S1**. Protein sequences of CHO-expressed Spike truncations. Signal sequences are highlighted in gray, and potential N-linked glycosylation sites are highlighted in magenta. Percentages indicate percent coverage of full-length Spike construct, including signal sequences and 6x His tags.

| Truncation | Amino acid sequence, including secretion signal and 6x His tag |
|---|---|
| T1 (23.5%) | MFVFLVLLPLVSSQNFRVQPTESIVRFPNITNLCPFGEVFNATRFASVYAWN RKRISNCVADYSVLYNSASFSTFKCYGVSPTKLNDLCFTNVYADSFVIRGDE VRQIAPGQTGKIADYNYKLPDDFTGCVIAWNSNNLDSKVGGNYNYLYRLFR KSNLKPFERDISTEIYQAGSTPCNGVEGFNCYFPLQSYGFQPTNGVGYQPY RVVVLSFELLHAPATVCGPKKSTNLVKNKCVNFNFNGLTGTGVLTESNKKFL PFQQFGRDIADTTDAVRDPQTLEILDITPCSHHHHHH |
| T2 (25.1%) | MFVFLVLLPLVSSQTAGAAAYYVGYLQPRTFLLKYNENGTITDAVDCALDPL SETKCTLKSFTVEKGIYQTSNFRVQPTESIVRFPNITNLCPFGEVFNATRFAS VYAWNRKRISNCVADYSVLYNSASFSTFKCYGVSPTKLNDLCFTNVYADSF VIRGDEVRQIAPGQTGKIADYNYKLPDDFTGCVIAWNSNNLDSKVGGNYNYL YRLFRKSNLKPFERDISTEIYQAGSTPCNGVEGFNCYFPLQSYGFQPTNGV GYQPYRVVVLSFELLHAPATVCGPKKSTNLVKNKCVNFNFNGLTGTGVLTE HHHHHH |
| T3 (28.1%) | MFVFLVLLPLVSSQTAGAAAYYVGYLQPRTFLLKYNENGTITDAVDCALDPL SETKCTLKSFTVEKGIYQTSNFRVQPTESIVRFPNITNLCPFGEVFNATRFAS VYAWNRKRISNCVADYSVLYNSASFSTFKCYGVSPTKLNDLCFTNVYADSF VIRGDEVRQIAPGQTGKIADYNYKLPDDFTGCVIAWNSNNLDSKVGGNYNYL YRLFRKSNLKPFERDISTEIYQAGSTPCNGVEGFNCYFPLQSYGFQPTNGV GYQPYRVVVLSFELLHAPATVCGPKKSTNLVKNKCVNFNFNGLTGTGVLTE SNKKFLPFQQFGRDIADTTDAVRDPQTLEILDITPCSHHHHHH |
| T4 (27.5%) | MFVFLVLLPLVSSQNFRVQPTESIVRFPNITNLCPFGEVFNATRFASVYAWN RKRISNCVADYSVLYNSASFSTFKCYGVSPTKLNDLCFTNVYADSFVIRGDE VRQIAPGQTGKIADYNYKLPDDFTGCVIAWNSNNLDSKVGGNYNYLYRLFR KSNLKPFERDISTEIYQAGSTPCNGVEGFNCYFPLQSYGFQPTNGVGYQPY RVVVLSFELLHAPATVCGPKKSTNLVKNKCVNFNFNGLTGTGVLTESNKKFL PFQQFGRDIADTTDAVRDPQTLEILDITPCSFGGVSVITPGTNTSNQVAVLYQ DVNCTEVPVAIHADQLTPTWRVYSTGSNVHHHHHH |

| | |
|---|---|
| T5 (28.3%) | MFVFLVLLPLVSSQGFSALEPLVDLPIGINITRFQTLLALHRSYLTPGDSSSG<br>WTAGAAAYYVGYLQPRTFLLKYNENGTITDAVDCALDPLSETKCTLKSFTVE<br>KGIYQTSNFRVQPTESIVRFPNITNLCPFGEVFNATRFASVYAWNRKRISNC<br>VADYSVLYNSASFSTFKCYGVSPTKLNDLCFTNVYADSFVIRGDEVRQIAPG<br>QTGKIADYNYKLPDDFTGCVIAWNSNNLDSKVGGNYNYLYRLFRKSNLKPF<br>ERDISTEIYQAGSTPCNGVEGFNCYFPLQSYGFQPTNGVGYQPYRVVVLSF<br>ELLHAPATVCGPKKSTNLVKNKCVNFNFNGLTGTGVLTEHHHHHH |
| T6 (35.4%) | MFVFLVLLPLVSSQGFSALEPLVDLPIGINITRFQTLLALHRSYLTPGDSSSG<br>WTAGAAAYYVGYLQPRTFLLKYNENGTITDAVDCALDPLSETKCTLKSFTVE<br>KGIYQTSNFRVQPTESIVRFPNITNLCPFGEVFNATRFASVYAWNRKRISNC<br>VADYSVLYNSASFSTFKCYGVSPTKLNDLCFTNVYADSFVIRGDEVRQIAPG<br>QTGKIADYNYKLPDDFTGCVIAWNSNNLDSKVGGNYNYLYRLFRKSNLKPF<br>ERDISTEIYQAGSTPCNGVEGFNCYFPLQSYGFQPTNGVGYQPYRVVVLSF<br>ELLHAPATVCGPKKSTNLVKNKCVNFNFNGLTGTGVLTESNKKFLPFQQFG<br>RDIADTTDAVRDPQTLEILDITPCSFGGVSVITPGTNTSNQVAVLYQDVNCTE<br>VPVAIHADQLTPTWRVYSTGSNVHHHHHH |
| T7 (35.5%) | MFVFLVLLPLVSSQNFRVQPTESIVRFPNITNLCPFGEVFNATRFASVYAWN<br>RKRISNCVADYSVLYNSASFSTFKCYGVSPTKLNDLCFTNVYADSFVIRGDE<br>VRQIAPGQTGKIADYNYKLPDDFTGCVIAWNSNNLDSKVGGNYNYLYRLFR<br>KSNLKPFERDISTEIYQAGSTPCNGVEGFNCYFPLQSYGFQPTNGVGYQPY<br>RVVVLSFELLHAPATVCGPKKSTNLVKNKCVNFNFNGLTGTGVLTESNKKFL<br>PFQQFGRDIADTTDAVRDPQTLEILDITPCSFGGVSVITPGTNTSNQVAVLYQ<br>DVNCTEVPVAIHADQLTPTWRVYSTGSNVFQTRAGCLIGAEHVNNSYECDIP<br>IGAGICASYQTQTNSPASVASQSIIAYTMSLGAENSVAYSNNSIAIPTNFTISV<br>TTEILPVSMTKTSVDCTMYICGDHHHHHH |
| T8 (36.8%) | MFVFLVLLPLVSSQKTQSLLIVNNATNVVIKVCEFQFCNDPFLGVYYHKNNKS<br>WMESEFRVYSSANNCTFEYVSQPFLMDLEGKQGNFKNLREFVFKNIDGYFK<br>IYSKHTPINLVRDLPQGFSALEPLVDLPIGINITRFQTLLALHRSYLTPGDSSS<br>GWTAGAAAYYVGYLQPRTFLLKYNENGTITDAVDCALDPLSETKCTLKSFTV<br>EKGIYQTSNFRVQPTESIVRFPNITNLCPFGEVFNATRFASVYAWNRKRISN<br>CVADYSVLYNSASFSTFKCYGVSPTKLNDLCFTNVYADSFVIRGDEVRQIAP<br>GQTGKIADYNYKLPDDFTGCVIAWNSNNLDSKVGGNYNYLYRLFRKSNLKP<br>FERDISTEIYQAGSTPCNGVEGFNCYFPLQSYGFQPTNGVGYQPYRVVVLS<br>FELLHAPATVCGPKKSTNLVKNKCVNFNFNGLTGTGVLTEHHHHHH |
| S1 (54.7%) | MFVFLVLLPLVSSQCVNLTTRTQLPPAYTNSFTRGVYYPDKVFRSSVLHSTQ<br>DLFLPFFSNVTWFHAIHVSGTNGTKRFDNPVLPFNDGVYFASTEKSNIIRGWI<br>FGTTLDSKTQSLLIVNNATNVVIKVCEFQFCNDPFLGVYYHKNNKSWMESEF<br>RVYSSANNCTFEYVSQPFLMDLEGKQGNFKNLREFVFKNIDGYFKIYSKHTP<br>INLVRDLPQGFSALEPLVDLPIGINITRFQTLLALHRSYLTPGDSSSGWTAGA<br>AAYYVGYLQPRTFLLKYNENGTITDAVDCALDPLSETKCTLKSFTVEKGIYQT<br>SNFRVQPTESIVRFPNITNLCPFGEVFNATRFASVYAWNRKRISNCVADYSV<br>LYNSASFSTFKCYGVSPTKLNDLCFTNVYADSFVIRGDEVRQIAPGQTGKIA<br>DYNYKLPDDFTGCVIAWNSNNLDSKVGGNYNYLYRLFRKSNLKPFERDISTE<br>IYQAGSTPCNGVEGFNCYFPLQSYGFQPTNGVGYQPYRVVVLSFELLHAPA<br>TVCGPKKSTNLVKNKCVNFNFNGLTGTGVLTESNKKFLPFQQFGRDIADTT<br>DAVRDPQTLEILDITPCSFGGVSVITPGTNTSNQVAVLYQDVNCTEVPVAIHA<br>DQLTPTWRVYSTGSNVFQTRAGCLIGAEHVNNSYECDIPIGAGICASYQTQT<br>NSPAHHHHHH |

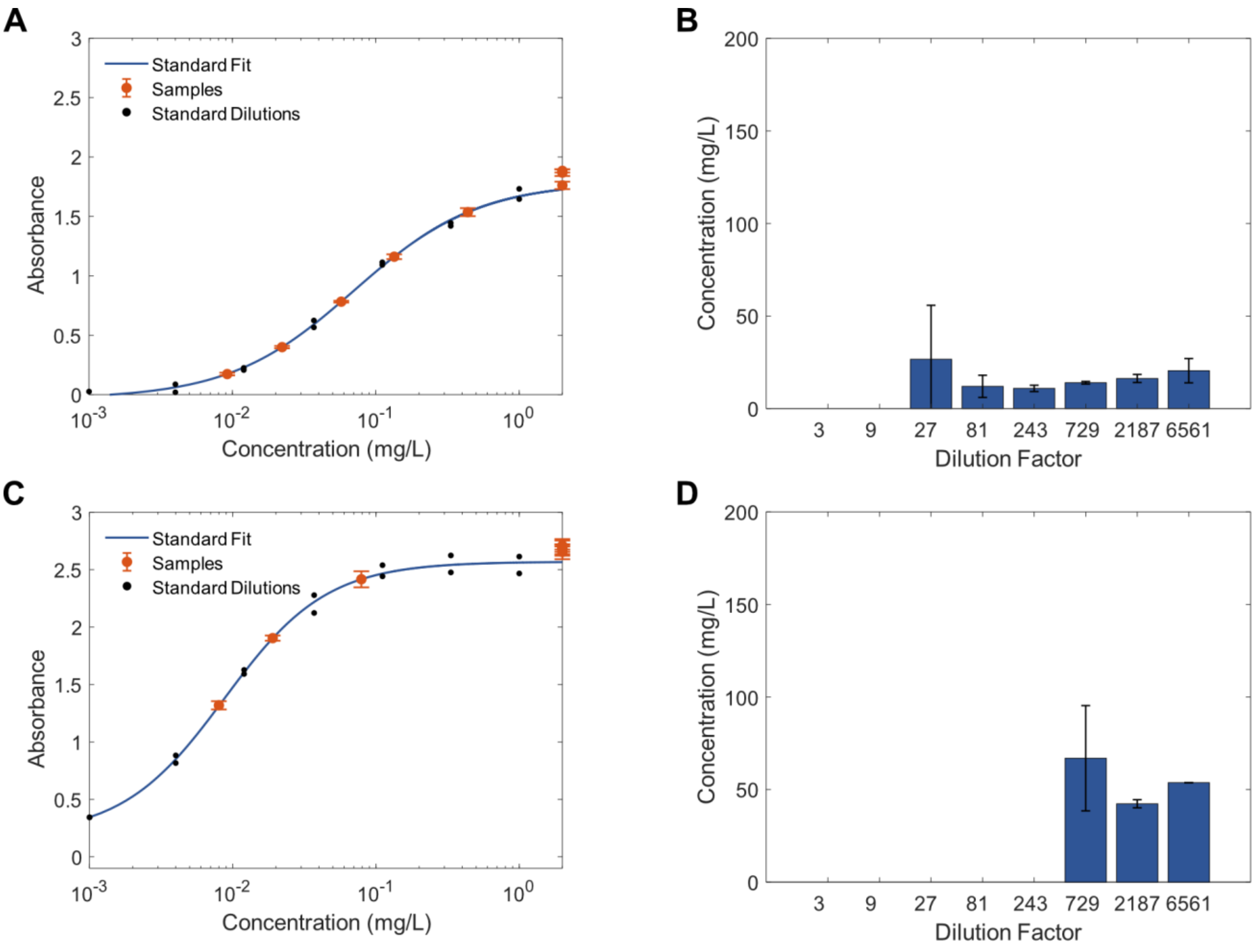


**Figure S1.** Measurement of crude titers for Spike and RBD via sandwich ELISA. (**A**) Dilutions of crude Spike plotted against the Spike standard curve. (**B**) Back-calculated concentrations of Spike crude titers. (**C**) Dilutions of crude RBD plotted against the RBD standard curve. (**D**) Back-calculated concentrations of RBD crude titers. Error bars represent ± SD of technical triplicates.

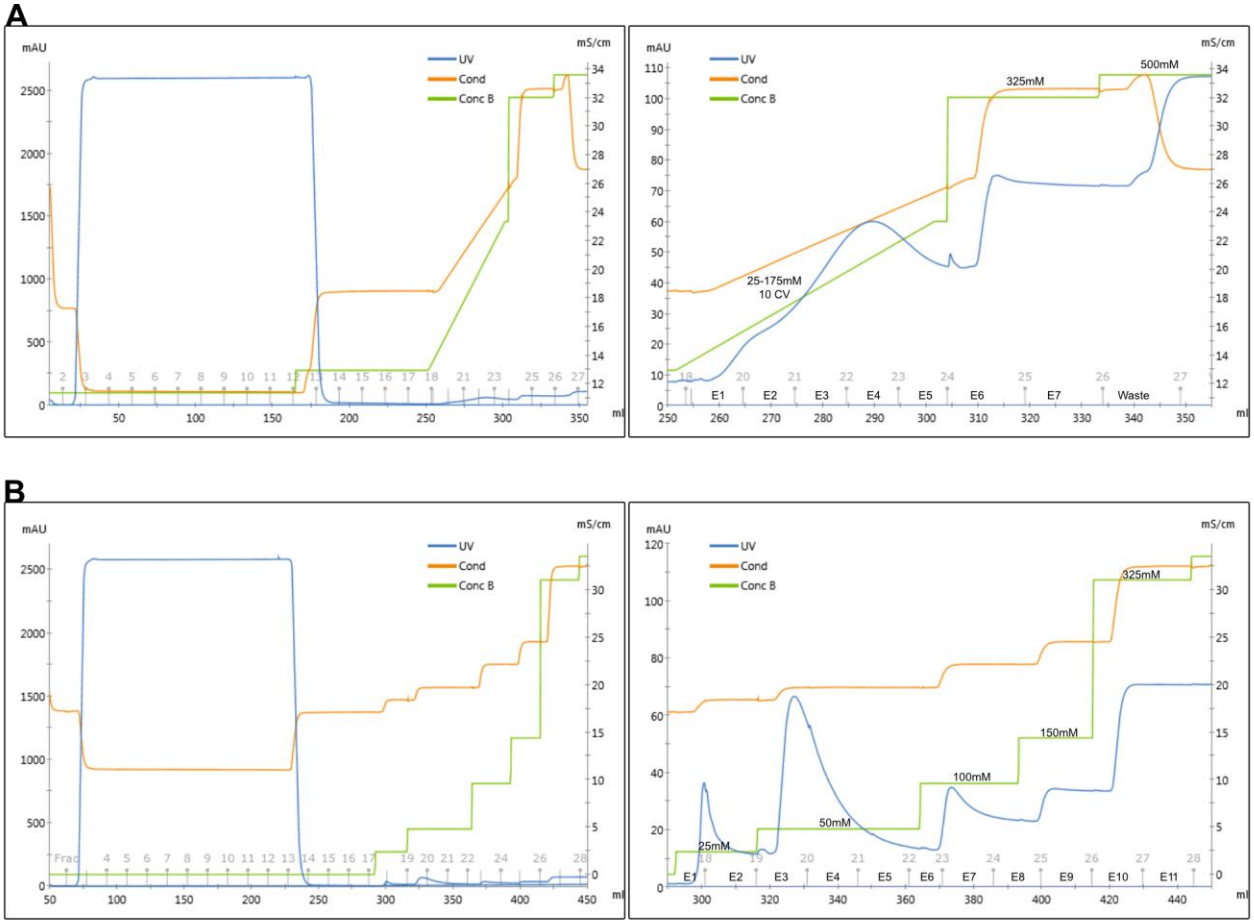


**Figure S2.** Spike and RBD chromatograms plotted over volume flowed through the system. (**A**) Chromatogram of entire Spike purification (left) and of zoomed in elution fractions (right). 150 mL of sample was loaded and washed with PBS containing 25 mM imidazole. A continuous gradient was applied from 25 mM-175 mM imidazole over 10 CV to elute Spike. Fractions E3 and E4 were collected. (**B**) Chromatogram of entire RBD purification (left) and of zoomed in elution fractions (right). 150 mL of sample was loaded and washed for 10 CV with PBS. A step gradient was applied for elution with 6 CV steps at 25 mM, 50 mM, 100 mM, 150 mM, and 325 mM imidazole. Fractions E3 and E4 were collected. Y-axis on the left is UV absorbance and Y-axis on the right is conductivity.

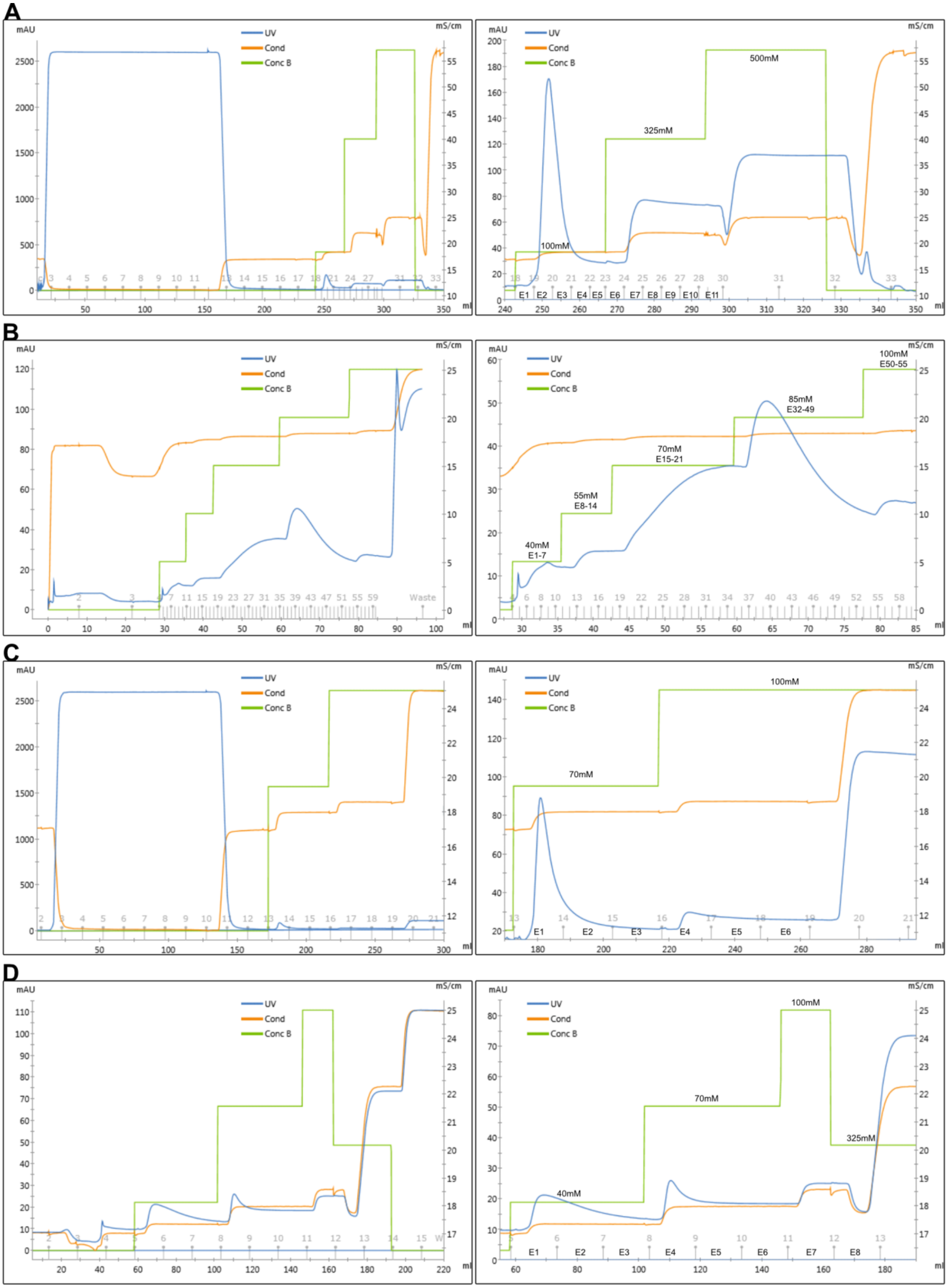

A
mAU
mS/cm
UV
Cond
Conc B
ml
100mM
325mM
500mM
E1 E2 E3 E4 E5 E6 E7 E8 E9 E10 E11
B
40mM
E1-7
55mM
E8-14
70mM
E15-21
85mM
E32-49
100mM
E50-55
Waste
C
70mM
100mM
E1 E2 E3 E4 E5 E6
D
40mM
70mM
100mM
325mM
E1 E2 E3 E4 E5 E6 E7 E8

**Figure S3.** T1 and T4 purification chromatograms. (**A**) Chromatogram of entire T1 purification (left) and of zoomed in elution fractions (right). First, 150 mL of sample was loaded and washed with PBS containing 25 mM imidazole. Step gradients were applied at 100 mM and 325 mM imidazole to elute T1. (**B**) Fractions E2 and E3 from the first purification were combined, dialyzed and re-purified using a step gradient with steps at 40 mM, 55 mM, 70 mM, 85 mM, and 100 mM imidazole. Fractions eluting from 40-85 mM imidazole were combined. (**C**) Chromatogram of entire T4 purification (left) and of zoomed in elution fractions (right). First, 120 mL of sample was loaded, and the resin was washed with PBS containing 25 mM imidazole. A step gradient at 70 mM and 100 mM imidazole was used to elute T4. (**D**) T4 from the first purification was dialyzed and re-purified using step gradients at 40 mM, 70 mM, 100 mM, 325 mM, and 500 mM imidazole. 40 mM imidazole fractions and the first 70 mM imidazole fraction were combined and used for further characterization.

**A**

```
Ref_T1_Seq      NFRVQPTESIVRFPNITNLCPFGEVFNATRFASVYAWNRKRISNCVADYSVLYNSASFST
T1_Coverage     -FRVQPTESIVR------------------FASVYAWNRKRISNCVADYSVLYNSASFST
                 ***********                  ******************************

Ref_T1_Seq      FKCYGVSPTKLNDLCFTNVYADSFVIRGDEVRQIAPGQTGKIADYNYKLPDDFTGCVIAW
T1_Coverage     FKCYGVSPTKLNDLCFTNVYADSFVIRGDEVRQIAPGQTGKIADYNYKLPDDFTGCVIAW
                ************************************************************

Ref_T1_Seq      NSNNLDSKVGGNYNYLYRLFRKSNLKPFERDISTEIYQAGSTPCNGVEGFNCYFPLQSYG
T1_Coverage     NSNNLDSKVGGNYNYLYR---KSNLKPFERDISTEIYQAGSTPCNGVEGFNCYFPLQSYG
                ******************   ***************************************

Ref_T1_Seq      FQPTNGVGYQPYRVVVLSFELLHAPATVCGPKKSTNLVKNKCVNFNFNGLTGTGVLTESN
T1_Coverage     FQPTNGVGYQPYRVVVLSFELLHAPATVCGPKKSTNLVKNKCVNFNFNGLTGTGVLTESN
                ************************************************************

Ref_T1_Seq      KKFLPFQQFGRDIADTTDAVRDPQTLEILDITPCSHHHHHH
T1_Coverage     KKFLPFQQFGRDIADTTDAVRDPQTLEILDITPCSHHHHHH
                *****************************************
```

**B**

```
Ref_T4_Seq      NFRVQPTESIVRFPNITNLCPFGEVFNATRFASVYAWNRKRISNCVADYSVLYNSASFST
T4_Top_Coverage --RVQPTESIVR------------------FASVYAWNRKRISNCVADYSVLYNSASFST
                  **********                  ******************************

Ref_T4_Seq      FKCYGVSPTKLNDLCFTNVYADSFVIRGDEVRQIAPGQTGKIADYNYKLPDDFTGCVIAW
T4_Top_Coverage FKCYGVSPTKLNDLCFTNVYADSFVIRGDEVRQIAPGQTGKIADYNYKLPDDFTGCVIAW
                ************************************************************

Ref_T4_Seq      NSNNLDSKVGGNYNYLYRLFRKSNLKPFERDISTEIYQAGSTPCNGVEGFNCYFPLQSYG
T4_Top_Coverage NSNNLDSKVGGNYNYLYR---KSNLKPFERDISTEIYQAGSTPCNGVEGFNCYFPLQSYG
                ******************   ***************************************

Ref_T4_Seq      FQPTNGVGYQPYRVVVLSFELLHAPATVCGPKKSTNLVKNKCVNFNFNGLTGTGVLTESN
T4_Top_Coverage FQPTNGVGYQPYRVVVLSFELLHAPATVCGPKKSTNLVKNKCVNFNFNGLTGTGVLTESN
                ************************************************************

Ref_T4_Seq      KKFLPFQQFGRDIADTTDAVRDPQTLEILDITPCSFGGVSVITPGTNTSNQVAVLYQDVN
T4_Top_Coverage KKFLPFQQFGRDIADTTDAVRDPQTLEILDITPCS-------------------------
                ***********************************

Ref_T4_Seq      CTEVPVAIHADQLTPTWRVYSTGSNVHHHHHH
T4_Top_Coverage --------------------------HHHHHH
                                          ******
```

**C**

```
Ref_T4_Seq      NFRVQPTESIVRFPNITNLCPFGEVFNATRFASVYAWNRKRISNCVADYSVLYNSASFST
T4_Bottom_Cover -FRVQPTESIVR------------------FASVYAWNRKRISNCVADYSVLYNSASFST
                 ***********                  ******************************

Ref_T4_Seq      FKCYGVSPTKLNDLCFTNVYADSFVIRGDEVRQIAPGQTGKIADYNYKLPDDFTGCVIAW
T4_Bottom_Cover FKCYGVSPTKLNDLCFTNVYADSFVIRGDEVRQIAPGQTGKIADYNYKLPDDFTGCVIAW
                ************************************************************

Ref_T4_Seq      NSNNLDSKVGGNYNYLYRLFRKSNLKPFERDISTEIYQAGSTPCNGVEGFNCYFPLQSYG
T4_Bottom_Cover NSNNLDSKVGGNYNYLYR---KSNLKPFERDISTEIYQAGSTPCNGVEGFNCYFPLQSYG
                ******************   ***************************************

Ref_T4_Seq      FQPTNGVGYQPYRVVVLSFELLHAPATVCGPKKSTNLVKNKCVNFNFNGLTGTGVLTESN
T4_Bottom_Cover FQPTNGVGYQPYRVVVLSFELLHAPATVCGPKK--------CVNFNFNGLTGTGVLTESN
                *********************************        *******************

Ref_T4_Seq      KKFLPFQQFGRDIADTTDAVRDPQTLEILDITPCSFGGVSVITPGTNTSNQVAVLYQDVN
T4_Bottom_Cover KKFLPFQQFGRDIADTTDAVRDPQTLEILDITPCS-------------------------
                ***********************************

Ref_T4_Seq      CTEVPVAIHADQLTPTWRVYSTGSNVHHHHHH
T4_Bottom_Cover --------------------------HHHHHH
                                          ******
```

**Figure S4.** Shotgun proteomics on T1 and T4. Coverage of (**A**) T1, (**B**) T4 top band, and (**C**) T4 bottom band against full sequences. Tandem mass spectra were extracted by MS Convert (ProteoWizard). Charge state deconvolution and deisotoping were not performed. All MS/MS samples were analyzed using X! Tandem (The GPM, thegpm.org; version X! Tandem Alanine (2017.2.1.4)). X! Tandem was set up to search the Uniprot human database and known T1 and T4 sequences assuming the digestion enzyme trypsin. X! Tandem was searched with a fragment ion mass tolerance of 20 PPM and a parent ion tolerance of 20 PPM. Carbamidomethyl of cysteine and selenocysteine was specified in X! Tandem as a fixed modification. Glu->pyro-Glu of the n-terminus, ammonia-loss of the n-terminus, gln->pyro-Glu of the n-terminus, deamidated of asparagine and glutamine, oxidation of methionine and tryptophan and dioxidation of methionine and tryptophan were specified in X! Tandem as variable modifications. Scaffold (version Scaffold_4.9.0, Proteome Software Inc., Portland, OR) was used to validate MS/MS based peptide and protein identifications. Peptide identifications were accepted if they could be established at greater than 98.0% probability by the Scaffold Local FDR algorithm. Peptide identifications were also required to exceed specific database search engine thresholds. Protein identifications were accepted if they could be established at greater than 5.0% probability to achieve an FDR less than 5.0% and contained at least 2identified peptides. Protein probabilities were assigned by the Protein Prophet algorithm (Nesvizhskii et al., 2003) Proteins that contained similar peptides and could not be differentiated based on MS/MS analysis alone were grouped to satisfy the principles of parsimony. Proteins sharing significant peptide evidence were grouped into clusters.

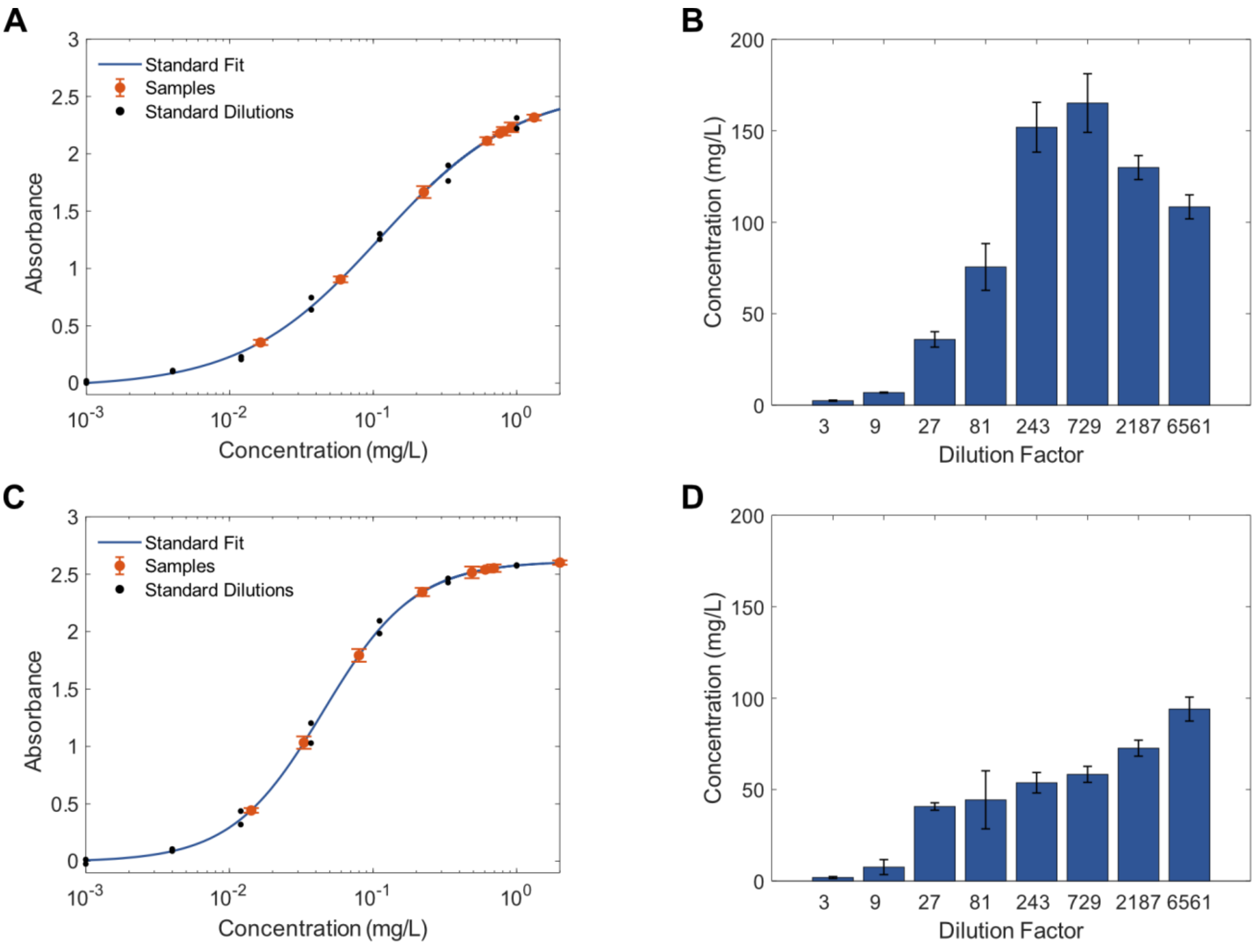


**Figure S5.** Measurement of crude titers for T1 and T4 via sandwich ELISA. (**A**) Dilutions of crude T1 plotted against the T1 standard curve. (**B**) Back-calculated concentrations of T1 crude titers. (**C**) Dilutions of crude T4 plotted against the T4 standard curve. (**D**) Back-calculated concentrations of T4 crude titers. Error bars represent ± SD of technical triplicates.

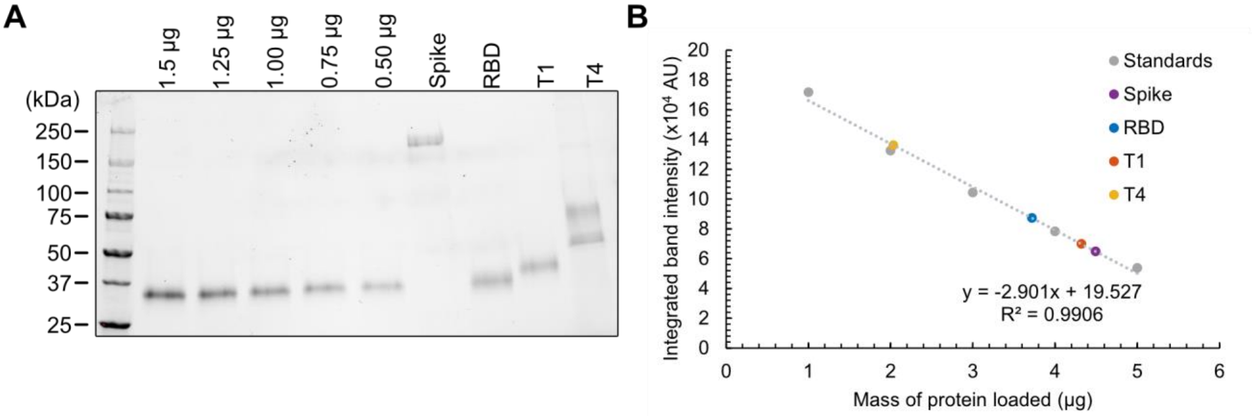


**Figure S6.** Quantification of purified proteins. (**A**) SDS-PAGE and (**B**) quantification for purified proteins and serial dilutions of RBD obtained from BEI Resources. A standard curve was prepared using a linear fit to serial dilutions of the standard protein.

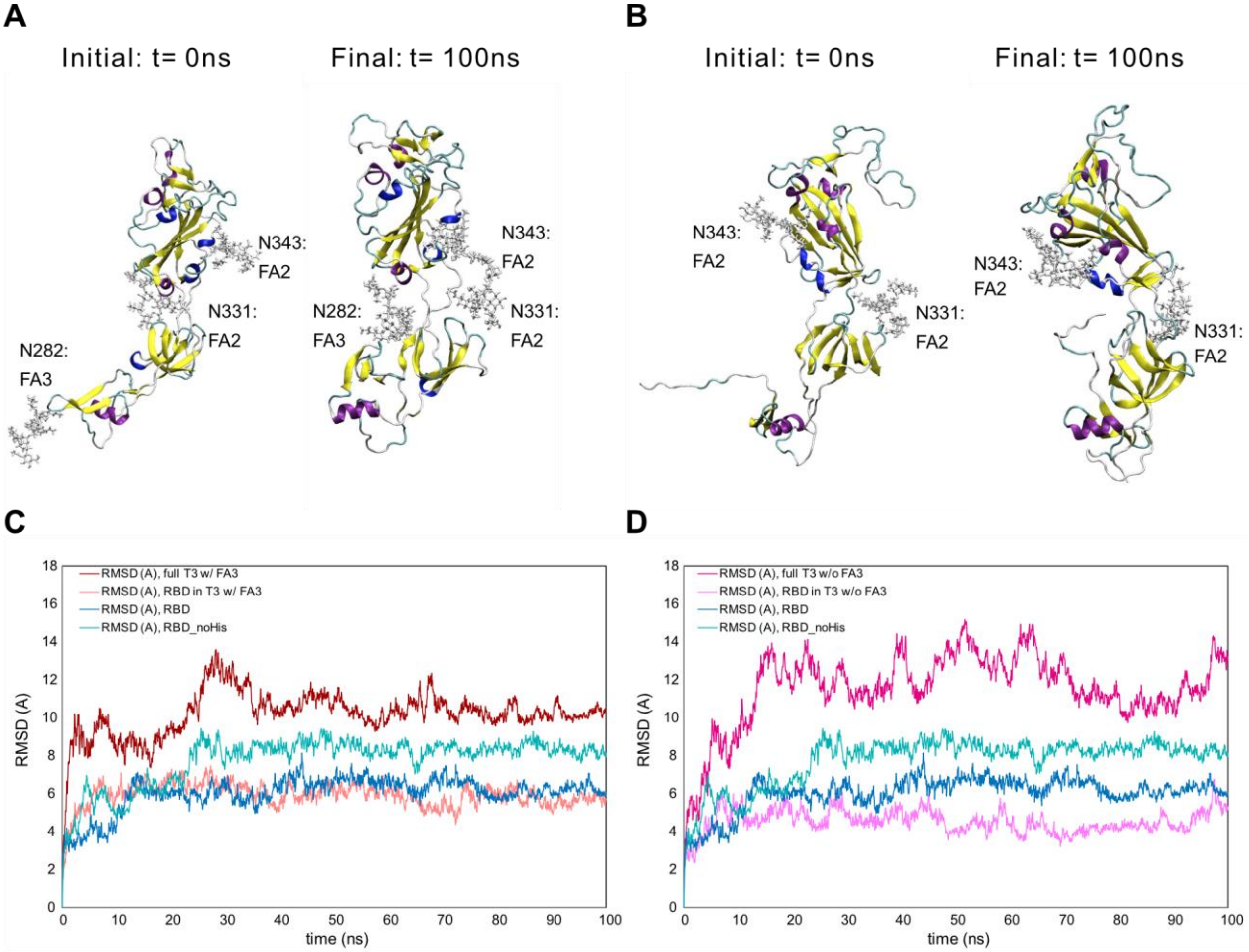


**Figure S7.** Molecular dynamics structural stability snapshots and analysis of T3. MD snapshots are visualized for (**A**) T3 with FA3 glycan and (**B**) T3 without FA3 glycan. (**C**) Backbone RMSD profiles of full T3 and T3 RBD subdomain with FA3 glycan. (D) Backbone RMSD profiles of full T3 and T3 RBD subdomain without FA3 glycan. Profiles of (C) and (D) include RBD with and without the 6x His tag referenced to initial configurations for comparison.

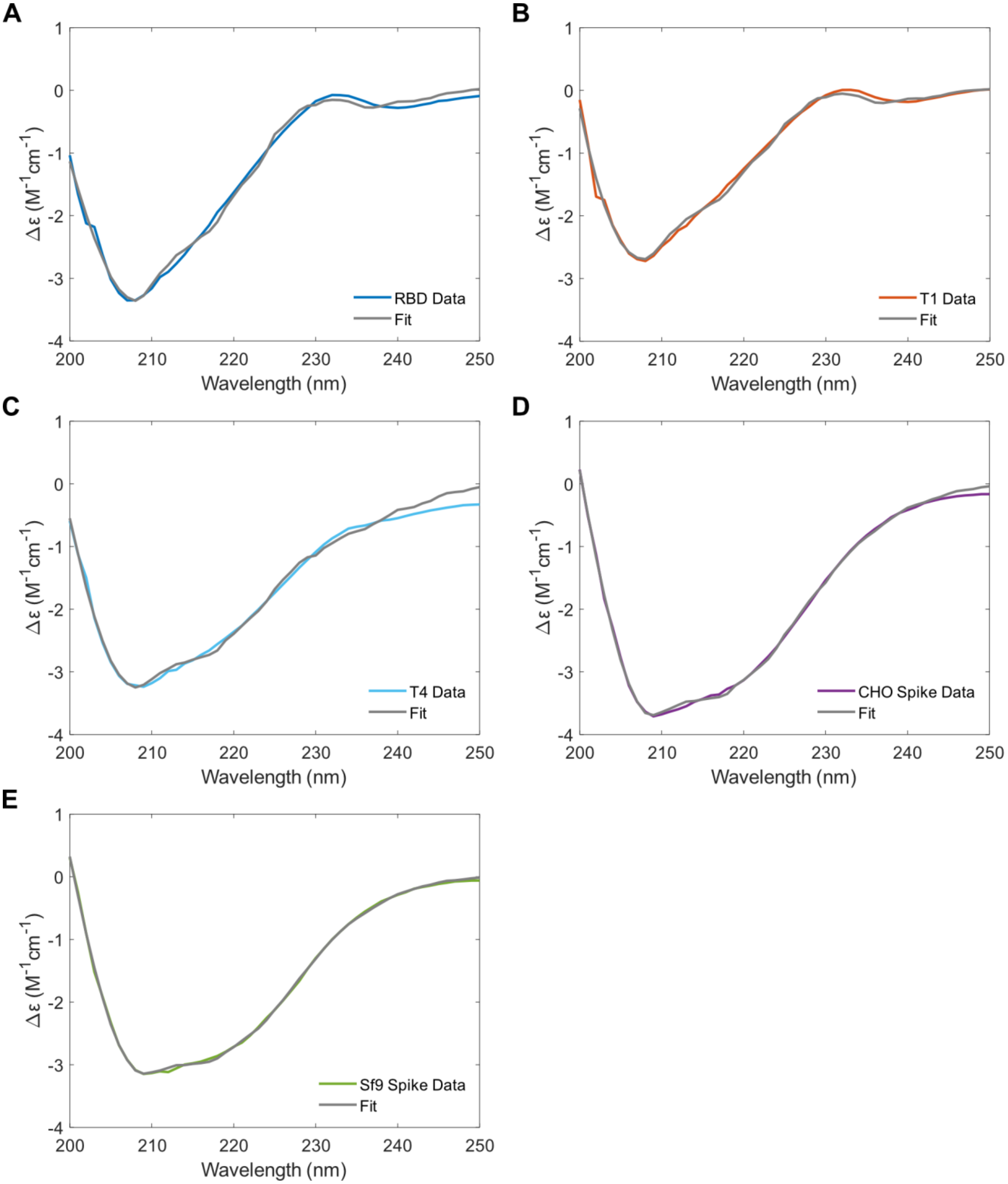


**Figure S8.** Raw spectral data on proteins analyzed via circular dichroism. Δε is plotted against wavelength for (**A**) RBD, (**B**) T1, (**C**) T4, (**D**) CHO Spike, and (**E**) Sf9 Spike.
